# bHLH11 inhibits bHLH IVc proteins by recruiting the TOPLESS/TOPLESS-RELATED corepressors in Arabidopsis

**DOI:** 10.1101/2020.04.09.035097

**Authors:** Yang Li, Rihua Lei, Mengna Pu, Yuerong Cai, Chengkai Lu, Zhifang Li, Gang Liang

## Abstract

Iron (Fe) homeostasis is essential for plant growth and development. Many transcription factors play pivotal roles in the maintenance of Fe homeostasis. bHLH11 was identified as a negative transcription factor regulating Fe homeostasis, however, the underlying molecular mechanism remains elusive. We generated two loss-of-function *bhlh11* mutants which display the enhanced sensitivity to Fe excess, the increased Fe accumulation and the elevated expression of Fe deficiency responsive genes. bHLH11 protein, localized in both the cytoplasm and nucleus, decreases in response to Fe deficiency. Coexpression assays indicate that bHLH IVc transcription factors (TFs) (bHLH34, bHLH104, bHLH105, and bHLH115) facilitate the nuclear accumulation of bHLH11 protein. Further analysis indicates that bHLH11 represses the transactivity of bHLH IVc TFs towards bHLH Ib genes (*bHLH38, bHLH39, bHLH100*, and *bHLH101*). bHLH11 contains two EAR motifs which are responsible for the repression function by recruiting the TOPLESS/TOPLESS-RELATED (TPL/TPRs) corepressors. Correspondingly, the expression of Fe uptake genes increases in the *tpr1 tpr4 tpl* mutant. Moreover, genetic analysis reveals that bHLH11 has functions independent of FIT. This study provides insights into the complicate Fe homeostasis signaling network.

**One-sentence summary:** bHLH IVc proteins promote the bHLH11 protein accumulation in the nucleus where bHLH11 inhibits the transcriptional activation ability of bHLH IVc via its EAR motifs recruiting the TOPLESS/TOPLESS-RELATED corepressors.

## Introduction

Iron (Fe) is an indispensable microelement for plant growth and development. Plants acquire Fe from the soil, which has low concentration of Fe available, especially in alkaline environments (Jeong and Guerinot, 2009). As about one-third of the world’s cultivated land is calcareous (alkaline), iron deficiency is common for plants (Grotz and Guerinot, 2006). Fe functions in many physiological processes, such as photosynthesis, respiration, hormone biosynthesis, and nitrogen fixation. Fe deficiency causes symptoms including delayed growth and leaf chlorosis and can affect the yield and nutritional quality of crops (Kobayashi and Nishizawa, 2014). Although Fe is required for plant growth and development, Fe excess can be toxic to plants because Fe can cause the production of reactive oxygen radicals that are harmful to plant cells (Quinet et al., 2012). Therefore, maintaining Fe homeostasis in plant cells is crucial for their normal growth and development.

Plants have evolved a set of molecular mechanisms for iron absorption, transport, distribution, and storage that ensure appropriate Fe concentrations in cells under low Fe availability. Dicotyledonous and non-gramineous monocotyledonous plants take up Fe using a reduction strategy (strategy I). In *Arabidopsis thaliana*, this strategy involves rhizosphere acidification, ferric iron reduction, and ferrous iron transport. H^+^-ATPases such as the P-type ATPase AHA2/AHA7 release protons into the soil, which improves the solubility of Fe in the soil (Santi and Schmidt, 2009; Kobayashi and Nishizawa, 2012). Then, the root surface Fe chelate reductase FERRIC REDUCTION OXIDASE2 (FRO2) catalyzes the reduction of Fe^3+^ to Fe^2+^ (Robinson et al., 1999). IRON-REGULATED TRANSPORTER1 (IRT1) transports Fe^2+^ into roots (Henriques et al., 2002; Varotto et al., 2002; Vert et al., 2002). By contrast, gramineous plants employ a chelation strategy (strategy II) in which high-affinity Fe chelators of the mugineic acid family, also known as phytosiderophores, are secreted into the rhizosphere and facilitate the uptake of the Fe^3+^-phytosiderophore complex. Recent studies suggest that secretion of Fe-chelating compounds is also important for the survival of non-gramineous plants such as Arabidopsis in alkaline soil (Rodríguez-Celm et al., 2013; Schmidt et al., 2014; Fourcroy et al., 2014, 2016; Siwinska et al., 2018; Tsai et al., 2018).

To maintain Fe homeostasis, plants must sense the environmental Fe concentration and fine-tune the expression of Fe uptake-associated genes accordingly. BRUTUS (BTS) interacts with the basic helix-loop-helix transcription factors bHLH105 and bHLH115 and promotes their degradation (Selote et al., 2015). IRON MAN (IMA), a class of peptides, interact with and inhibit BTS, facilitating the accumulation of bHLH105 and bHLH115 (Grillet et al., 2018; Li et al., 2021). bHLH105 and bHLH115 belong to the bHLH IVc group, which contains four members. The other two members are bHLH34 and bHLH104. These four members regulate the expression of *FER-LIKE IRON-DEFICIENCY-INDUCED TRANSCRIPTION FACTOR (FIT*), bHLH Ib genes (*bHLH38, bHLH39, bHLH100*, and *bHLH101*) and *POPEYE* (*PYE*) (Zhang et al., 2015; Li et al., 2016; Liang et al., 2017). bHLH121 interacts with bHLH IVc proteins and is required for the maintenance of Fe homeostasis (Kim et al., 2019; Gao et al., 2020; Lei et al., 2020). Downstream of bHLH IVc and bHLH121, FIT interacts with bHLH Ib TFs to promote the expression of Fe-uptake associated genes *IRT1* and *FRO2* (Yuan et al., 2008; Wang et al., 2013). In contrast, PYE and bHLH11 are negative regulators of Fe homeostasis (Long et al., 2010; Tanabe et al., 2019). Additionally, bHLH105 also functions as a negative regulator when it interacts with PYE (Tissot et al., 2019). There is also a similar Fe deficiency response signaling network in rice (Ogo et al., 2007; Kobayashi 2013, 2019; Zhang et al., 2017, 2020; Wang et al., 2020; Li et al., 2020).

The overexpression of *bHLH11* causes the dramatic decline of Fe uptake genes including *IRT1* and *FRO2*, and the severe Fe deficiency symptoms (Tanabe et al., 2019). However, the molecular mechanism by which bHLH11 regulates Fe homeostasis remains elusive. In the present study, we characterized the roles of bHLH11 in the maintenance of Fe homeostasis in Arabidopsis. bHLH11 is localized in both the cytoplasm and nucleus and is exclusively in the nucleus when bHLH IVc TFs are abundant. bHLH11 also interacts with and inhibits bHLH IVc TFs. bHLH11 exerts its transcriptional repression function by its two EAR motifs recruiting the transcriptional TOPLESS/TOPLESS-RELATED corepressors.

## Results

### Loss-of-function of *bHLH11* impairs Fe homeostasis

It was reported that the overexpression of *bHLH11* leads to the enhanced sensitivity to Fe deficiency (Tanabe et al., 2019). To further explore the molecular mechanism of bHLH11 regulating Fe homeostasis, we employed the CRISPR-Cas9 system to edit *bHLH11*. Two single guide RNAs were designed to specifically target exons 4 and 3 of *bHLH11* and respectively integrated into the binary vector with a Cas9 (Liang et al., 2016) which were then used for transformation of wild type plants. We identified two homozygous mutants (*bhlh11-1* and *bhlh11-2*), the former containing a 1-bp insertion in exon 4 and the latter containing a 2-bp deletion in exon 3 (Figure S1A), both of which caused a frameshift mutation (Figure S1B). Expression analysis indicated that bHLH11 mRNA levels did not change in these two mutants (Figure S1C). When grown on Fe0 (Fe free) or Fe100 (100 μM Fe^2+^) medium, no visible difference was observed between the *bhlh11* mutants and wild type (Figure 1A). By contrast, when grown on Fe300 (300 μM Fe^2+^) medium, the *bhlh11* mutants produced low shoot biomass (Figure 1A, B). The *bHLH11* driven by its native promoter rescued the sensitivity of *bhlh11-1* to Fe excess (Figure S1D). Fe content analysis suggested that the Fe concentration of *bhlh11* mutants was higher than that of the wild type (Figure 1C). These data suggest that the loss-of-function of *bHLH11* leads to the enhanced sensitivity to Fe excess.

**Figure 1.**
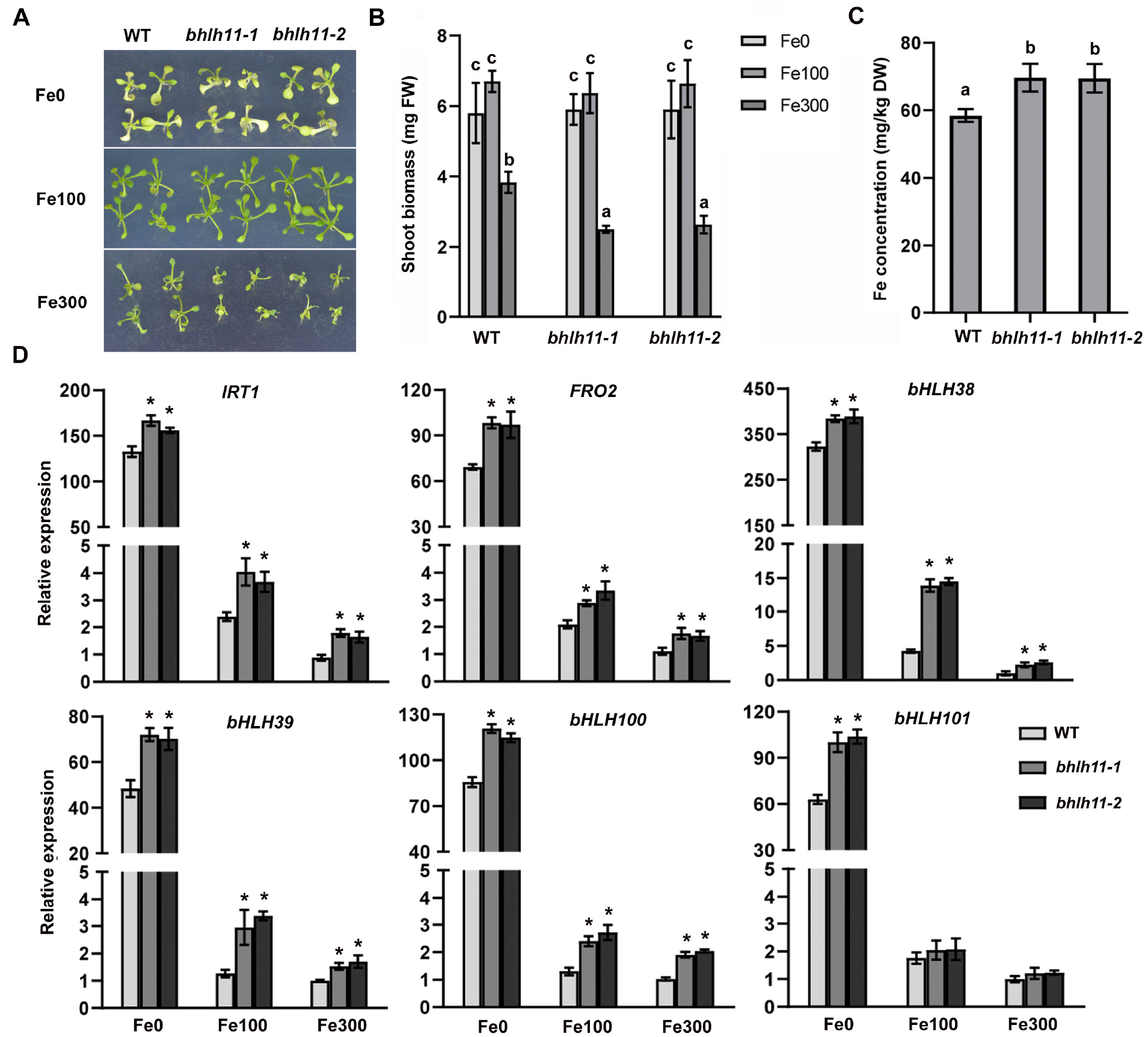
*bhlh11* mutants are sensitive to Fe excess. (A) Phenotypes of *bhlh11* mutants. Two-week-old seedlings grown on Fe0, Fe100 or Fe300 medium. (B) Shoot biomass of *bhlh11* mutants. Fresh weight of two-week-old shoots grown on Fe0, Fe100 or Fe300 medium. Three biological duplicates, each of which contains 15 plants, were analyzed. (C) Fe concentration of rosette leaves of three-week-old wild type and *bhlh11* plants grown in soil. (D) Expression of *IRT1, FRO2* and bHLH Ib genes. Four-day-old plants grown on Fe100 medium were transferred to Fe0, Fe100 or Fe300 medium for three days, and root samples were harvested and used for RNA extraction and RT-qPCR. (B-D)The different letters above each bar indicate statistically significant differences as determined by one-way ANOVA followed by Tukey’s multiple comparison test (P < 0.05).

To further investigate the effect of bHLH11 on the Fe signaling network, we examined the expression of several Fe homeostasis associated genes (Figure 1D). The expression of *IRT1, FRO2, bHLH38, bHLH39*, and *bHLH100* was higher in the *bhlh11* mutants than in the wild type regardless of Fe status. These data further support the negative regulation function of *bHLH11* in the Fe homeostasis.

### bHLH11 expression and subcellular localization

To investigate the response of *bHLH11* to Fe status, RT-qPCR was used to determine the expression of *bHLH11* in response to Fe status, showing that *bHLH11* mRNA increased in the roots with an increase of Fe concentration in the growth medium (Figure 2A), which is in consistence with the previous study (Tanabe et al., 2019). To examine the response of bHLH11 protein to Fe status, the *bHLH11* overexpression construct was introduced into wild type plants. In agreement with the previous report (Tanabe et al., 2019), *bHLH11* overexpression plants were more sensitive to Fe deficiency compared with wild type (Figure S2). One-week-old *bHLH11* overexpression plants grown on Fe100 medium were transferred to Fe0 or Fe300 medium, and root samples were harvested after 1, 2, and 3 days. Immunoblot analysis showed that bHLH11 increased with an increase in Fe concentration and decreased with a decrease in Fe concentration (Figure 2B).

**Figure 2.**
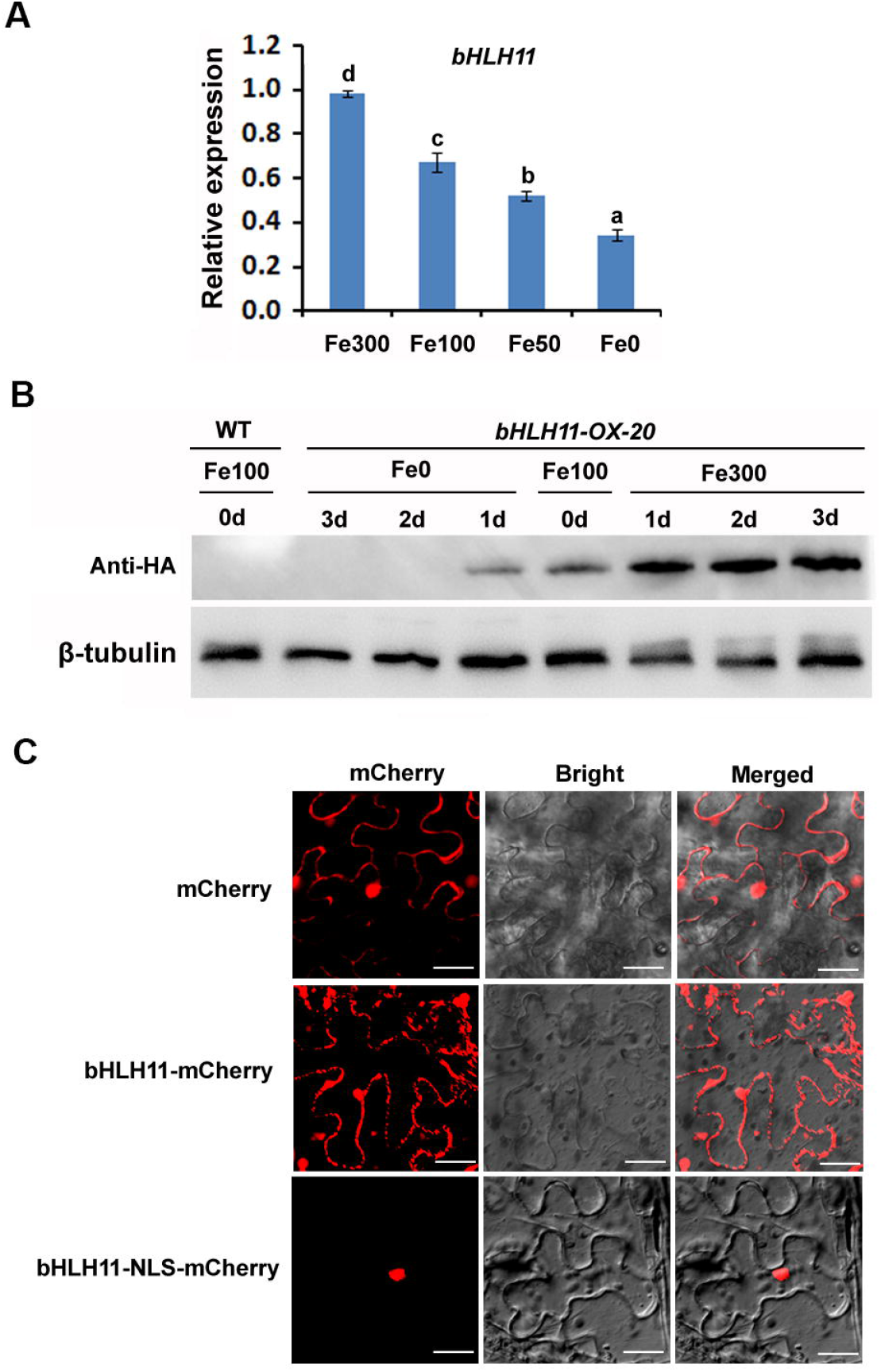
Response of bHLH11 to Fe status (A) RT-qPCR analysis of *bHLH11* expression. Four-day-old plants grown on Fe100 medium were shifted to Fe0, Fe50, Fe100 or F300 medium for 3 days. Roots were used for RNA extraction and RT-qPCR. The different letters above each bar indicate statistically significant differences as determined by one-way ANOVA followed by Tukey’s multiple comparison test (P < 0.05). (B) Degradation of bHLH11 in response to Fe deficiency. Seven-day-old wild type and *bHLH11-OX-20* seedlings grown on Fe100 medium were transferred to Fe0 or Fe300 medium, and root samples were harvested after 1, 2, and 3 days. anti-HA was used to detect HA-bHLH11. β-tubulin was used as a loading control. (C) Subcellular localization of bHLH11. The free mCherry, bHLH11-mCherry or bHLH11-NLS-mCherry were respectively expressed in *N. benthamiana* leaves.

Several Fe-homeostasis associated bHLH TFs were found outside the nucleus (Gratz et al., 2019; Trofimov et al., 2019; Lei et al., 2020; Wang et al., 2020; Liang et al., 2020). To determine the subcellular localization of bHLH11, we generated the *35S:bHLH11-mCherry* construct, in which the mCherry tag was fused in frame with the C end of bHLH11. When this construct was transiently expressed in tobacco leaves, mCherry was mainly observed in the cytoplasm and nucleus, which is very similar to that of free mCherry (Figure 2C). The cytoplasmic localization of bHLH11 was unexpected because transcription factors are known to function in the nucleus. Thus, we examined whether bHLH11 can be retained in the cytoplasm due to a lack of a nuclear localization signal (NLS). NLS prediction was conducted by cNLS Mapper with a cutoff score = 4 (Kosugi et al., 2009;http://nls-mapper.iab.keio.ac.jp/cgi-bin/NLS_Mapper_y.cgi). No NLS was found in bHLH11. When bHLH11 was fused with NLS-mCherry, which contains an NLS from the SV40 virus, bHLH11-NLS-mCherry was exclusively localized in the nucleus (Figure 2C). These data suggest that the lack of an NLS causes the cytoplasmic localization of bHLH11.

### bHLH11 interacts with bHLH IVc TFs in the nucleus

Considering that TFs usually function in the nucleus and an NLS allows bHLH11 to remain in the nucleus, we hypothesized that bHLH11 might be recruited to the nucleus by its nuclear-localized interaction partners. Recent studies revealed that bHLH121, the closest homolog of bHLH11, interacts with bHLH IVc TFs (Kim et al., 2019; Gao et al., 2020; Lei et al., 2020). Therefore, we employed the yeast two-hybrid system to test whether bHLH11 interacts with bHLH IVc TFs. The bHLH11 protein was fused with the GAL4 DNA binding domain in the pGBKT7 vector as the bait (BD-bHLH11). bHLH IVc TFs were cloned to the pGADT7 vector as the preys. As expected, bHLH11 interacts with all four bHLH IVc TFs in yeast (Figure 3A).

**Figure 3.**
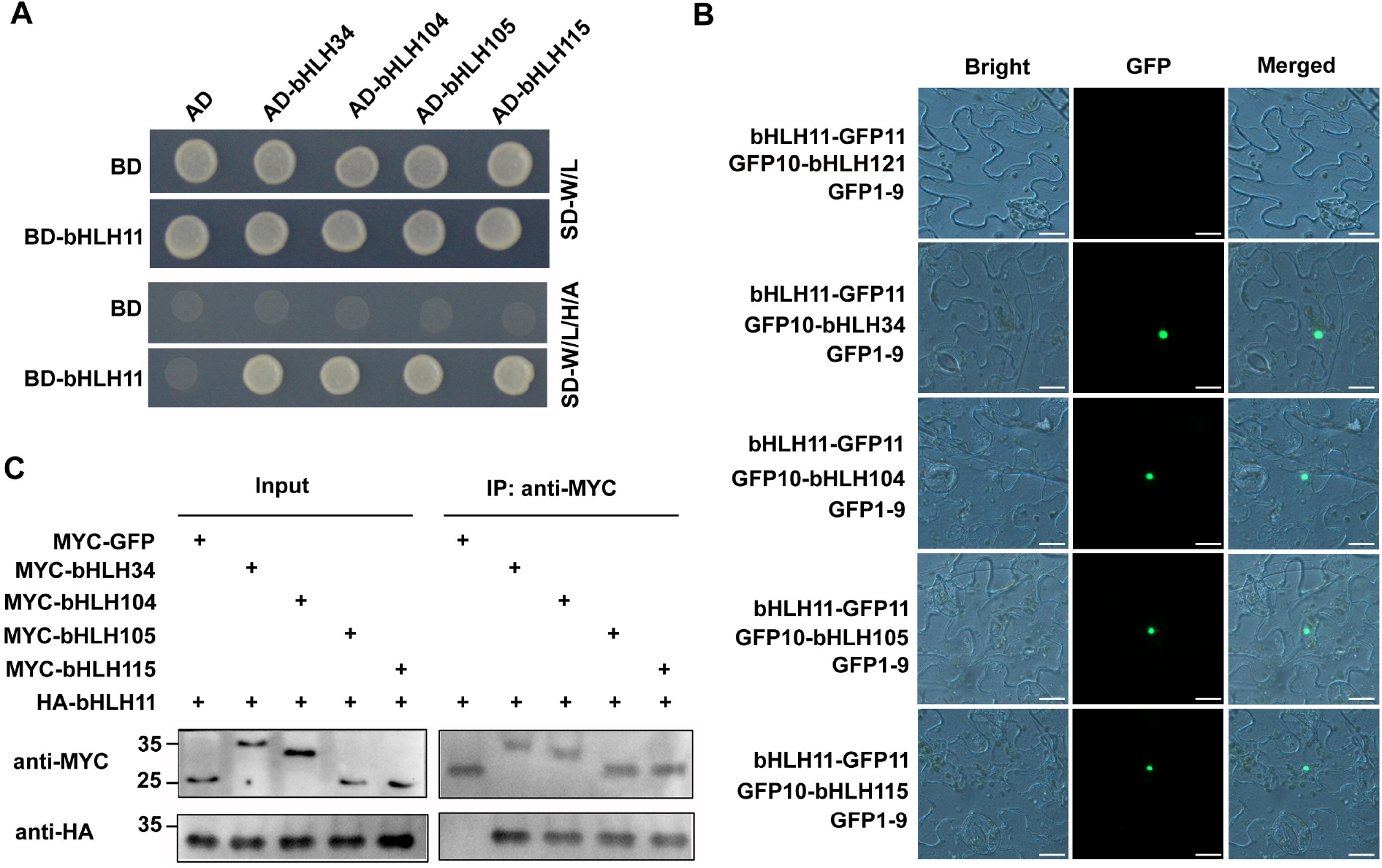
bHLH11 physically interacts with bHLH IVc TFs. (A) Yeast two-hybrid analysis of the interaction between bHLH11 and bHLH IVc TFs. Yeast cotransformed with different BD and AD plasmid combinations was spotted on synthetic dropout medium lacking Leu/Trp (SD–W/L) or Trp/Leu/His/Ade (SD–W/L/H/A). (B) Interaction of bHLH11 and bHLH IVc TFs in plant cells. Tripartite split-sfGFP complementation assays were performed. bHLH34, bHLH104, bHLH105, bHLH115, and bHLH121 were fused with GFP10, and bHLH11 was fused with GFP11. The constructs were introduced into agrobacterium respectively, and the indicated combinations were co-expressed in *N. benthamiana* leaves. (C) Co-IP analysis of the interaction between bHLH11 and bHLH IVc TFs. Total proteins from different combinations of HA-bHLH11 and MYC-GFP, MYC-bHLH34, MYC-bHLH104, MYC-bHLH105, or MYC-bHLH115 were immunoprecipitated with anti-MYC followed by immunoblotting with the indicated antibodies. MYC-GFP was used as a negative control. Protein molecular weight (in kD) is indicated to the left of the immunoblot.

To confirm that bHLH IVc TFs interact with bHLH11 in plant cells, we employed the tripartite split-GFP system (Liu et al., 2018). The GFP10 fragment was fused with bHLH IVc proteins in their N-end (GFP10-bHLH IVc) and the GFP11 was fused with bHLH11 in the C-end (bHLH11-GFP11). As a control, bHLH121 was fused with the GFP10. When GFP10-bHLH IVc and bHLH11-GFP11 were transiently co-expressed with GFP1-9 in tobacco leaves, the GFP signal was visible in the nucleus of transformed cells (Figure 3B). By contrast, the combination GFP10-bHLH121/bHLH11-GFP11/GFP1-9 did not result in visible GFP signal. The negative controls did not result in a GFP signal in the cells (Figure S3).

We next performed a coimmunoprecipitation (Co-IP) assay to confirm the interactions between bHLH IVc TFs and bHLH11 (Figure 3C). MYC tag-fused bHLH IVc TFs and HA tag-fused bHLH11 were transiently co-expressed in tobacco leaves. The total proteins were incubated with MYC antibody and A/G-agarose beads and then separated on SDS-PAGE for immunoblotting with HA antibody. Consistent with the results from the yeast two-hybrid and tripartite split-GFP assays, bHLH IVc and bHLH11 were present in the same protein complex. These data suggest that bHLH IVc TFs physically interact with bHLH11.

### bHLH IVc TFs affect the subcellular localization of bHLH11

Having confirmed that bHLH11 interacts with bHLH IVc TFs in the nucleus, we wondered whether the bHLH IVc TFs have an impact on the subcellular localization of bHLH11. When any one of these four GFP tagged proteins was respectively co-expressed with bHLH11-mCherry, bHLH11-mCherry accumulated exclusively in the nucleus (Figure 4A). By contrast, co-expression of the free GFP did not affect the subcellular localization of bHLH11-mCherry (Figure 4A).

**Figure 4.**
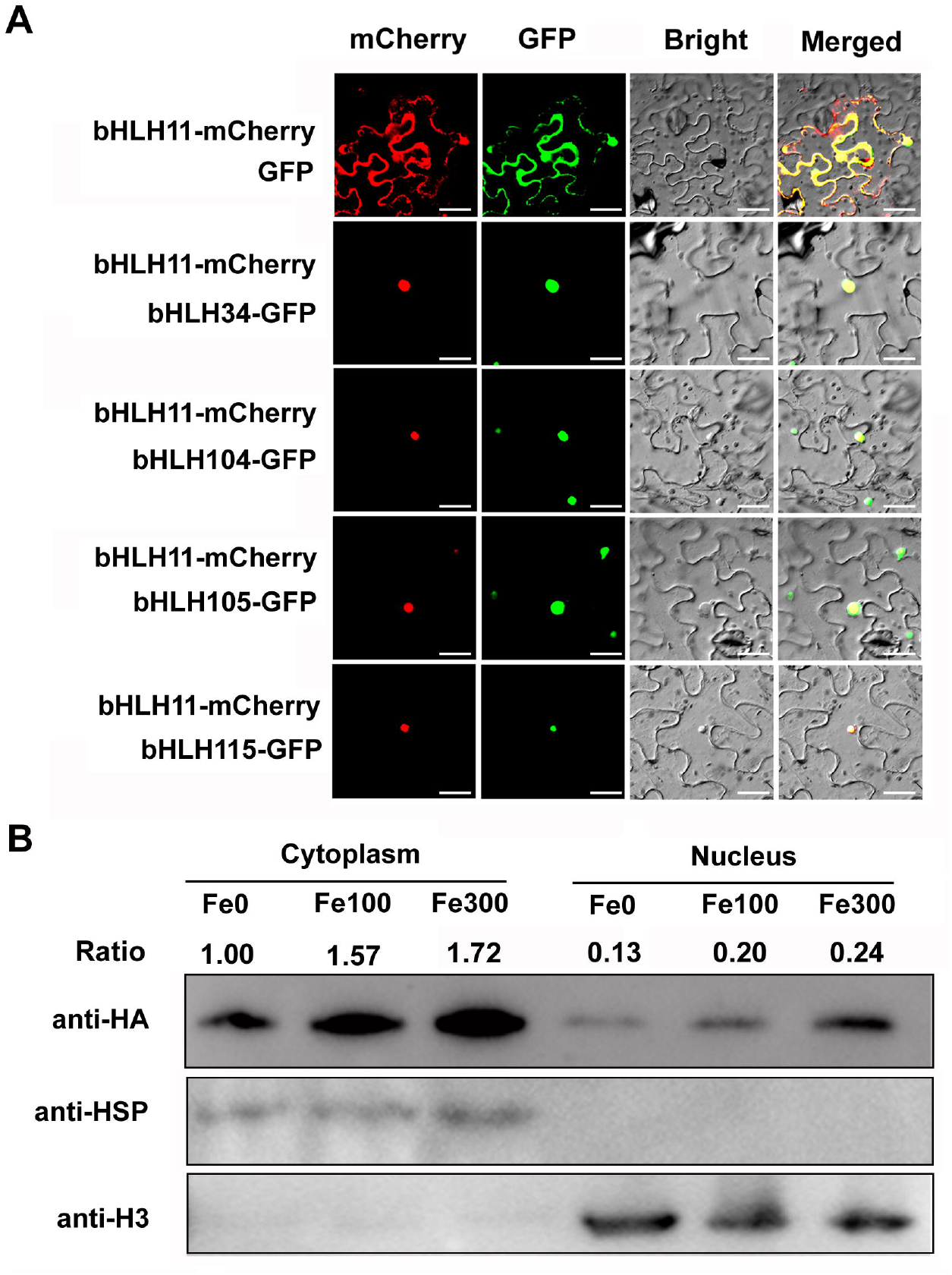
Change of bHLH11 subcellular localization. (A) Location of bHLH11 in the absence or presence of bHLH IVc. bHLH11-mCherry was co-expressed with bHLH IVc TFs. The combination of bHLH11-GFP and free mCherry was used as a negative control. Transient expression assays were performed in tobacco leaves. (B) Immunoblot analysis of bHLH11 protein distribution in the cytoplasm and nuclear fractions. Seven-day-old *bHLH11-OX-20* seedlings grown on Fe100 medium were transferred to Fe0, Fe100 or Fe300 medium. Root samples were harvested after 3 days, and cytoplasmic and nuclear proteins were extracted and subjected to immunoblot analysis with the indicated antibodies. Ratio indicates the relative protein abundance of HA-bHLH11.

To further confirm the distribution of bHLH11 in the cytoplasm and nucleus, we used immunoblot to measure the expression of the bHLH11 protein. As shown in Figure 4B, bHLH11 protein was detected both in the nucleus and cytoplasm, and both its nuclear and cytoplasmic counterparts were responsive to Fe status.

### bHLH11 antagonizes the transactivity of bHLH IVc TFs

The bHLH Ib genes are activated directly by the bHLH IVc TFs (Zhang et al., 2015; Li et al., 2016; Liang et al., 2017). Our expression analyses also suggested that the bHLH Ib genes were upregulated in the *bhlh11* mutants, implying that bHLH11 is a negative regulator of bHLH Ib genes. Because bHLH11 interacts with the bHLH IVc TFs, we proposed that bHLH11 could antagonize the functions of the bHLH IVc TFs. To confirm this hypothesis, transient expression assays were conducted in Arabidopsis protoplasts (Figure 5A). The reporter construct *ProbHLH38:LUC*, in which the LUC reporter was fused with the promoter of *bHLH38*, and different effectors in which the 35S promoter was used to drive GFP, bHLH11 or bHLH IVc, were used in the transient assays. Compared with GFP, bHLH IVc TFs activated the expression of *ProbHLH38:LUC*, whereas bHLH11 had no significant effect. When bHLH11 and bHLH IVc were co-expressed, the LUC/REN ratio declined significantly. These data suggest that bHLH11 inhibits the transactivity of bHLH IVc TFs towards *bHLH38*.

**Figure 5.**
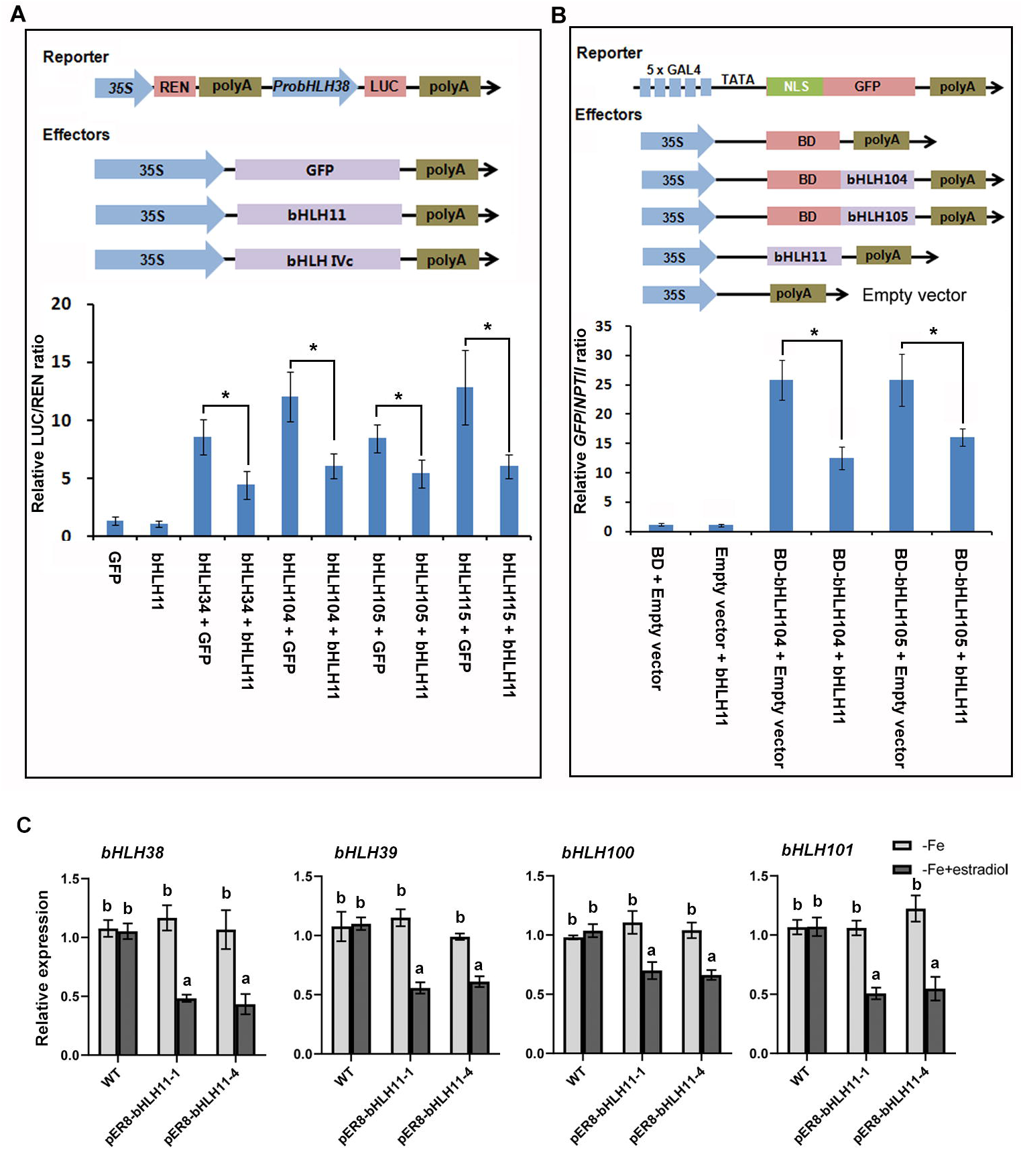
bHLH11 antagonizes the transcriptional activation ability of bHLH IVc TFs. (A) bHLH11 represses the functions of bHLH IVc TFs. Schematic diagram of the constructs transiently expressed in Arabidopsis protoplasts. The LUC/REN ratio represents the LUC activity relative to the internal control REN. The asterisk indicates a significant difference as determined by Student’s t Test. (B) bHLH11 inhibits the functions of bHLH IVc TFs by direct protein-protein interaction. The schematic diagram shows the constructs used in the transient expression assays in tobacco leaves. The abundance of *GFP* was normalized to that of *NPTII*. The asterisk indicates a significant difference as determined by Student’s t Test. (C) Expression of bHLH Ib genes in *pER8-bHLH11* plants. Seven-day-old plants grown on Fe0 medium were transferred to Fe0 medium with or without 4 μM estradiol for 6 h, and root samples were harvested and used for RNA extraction and RT-qPCR. The different letters above each bar indicate statistically significant differences as determined by one-way ANOVA followed by Tukey’s multiple comparison test (P < 0.05).

To further investigate whether bHLH11 inhibits the functions of bHLH IVc TFs by direct protein–protein interaction, we employed the *pGAL4* promoter. In the reporter construct, GFP fused with an NLS sequence was driven by *pGAL4* containing the minimal CaMV 35S promoter with five repeats of the GAL4 binding motif (Figure 5B). In the effectors, the DNA binding domain (BD) of GAL4 was fused in frame with either bHLH104 or bHLH105 and driven by the 35S promoter. Consistent with the fact that bHLH IVc TFs are transcriptional activators, the chimeric BD-bHLH104 or bHLH105 activated the expression of *GFP*. When bHLH11 was co-expressed with BD-bHLH104 or BD-bHLH105, the expression of *GFP* was significantly repressed. These data suggest that bHLH11 antagonizes the transcriptional activation ability of bHLH IVc TFs through direct protein interaction.

To further confirm the antagonistic role of bHLH11 to bHLH IVc TFs, we generated *bHLH11-OX/bHLH104-OX* plants by crossing *bHLH11-OX-20* with *bHLH104-OX*. Compared with *bHLH104-OX*, the tolerance of *bHLH11-OX/bHLH104-OX* to Fe deficiency was reduced (Figure S4). Taken together, our data suggest that bHLH11 can antagonizes the functions of bHLH IVc TFs.

It is reported that *bHLH11* overexpression causes the increased expression of bHLH Ib genes (Tanabe et al., 2019), which seems to be contrary to the results above that bHLH11 represses the expression of *bHLH38* (Figure 5A). We reasoned that the upregulation of bHLH Ib genes was not a direct result from the *bHLH11* overexpression, but caused by a secondary effect of the disrupted Fe homeostasis in *bHLH11-OX* plants. To avoid the secondary effect, we generated transgenic plants containing a *pER8-bHLH11* construct, in which the *HA-bHLH11* fusion gene was under the control of an inducible promoter, activated by estradiol. Under Fe deficient conditions, the *pER8-bHLH11* plants grew as well as the wild type, and the *bHLH11* transcript level was similar between the wild type and *pER8-bHLH11*. After treatment with estradiol, the *bHLH11* gene was overexpressed in the *pER8-bHLH11* plants (Figure S5A). As expected, the *pER8-bHLH11* plants displayed the enhanced sensitivity to Fe deficiency when gown on Fe0 + estradiol medium (Figure S5B). To examine the expression of Fe deficiency-responsive genes, seven-day-old plants grown on Fe0 medium were transferred to Fe0 medium with or without estradiol for 6 h and roots were isolated for analysis. We found that the expression of bHLH Ib genes was downregulated in the *pER8-bHLH11* plants after treatment with estradiol (Figure 5C). Taken together, our data suggest that bHLH11 antagonizes the transcriptional activation ability of bHLH IVc TFs to bHLH Ib genes.

### The repression function of bHLH11 requires its EAR motifs

Considering that bHLH11 negatively regulates the expression of Fe deficiency-responsive genes, we wanted to know how bHLH11 acts as a transcriptional repressor. There are two typical ethylene-responsive element binding factor-associated amphiphilic repression (EAR) motifs (LxLxL) in the C-terminal region of bHLH11 (Figure 6A). It is known that EAR motifs account for the repression function of many transcription factors (Kagale et al., 2010). To investigate whether the EAR motifs are required for the repression function of bHLH11, we conducted reporter–effector transient expression assays in which bHLH105 was used as an effector to activate *ProbHLH38-LUC* (Figure 6B). We compared the effects of GFP, bHLH11, bHLH11dm (a version containing two mutated EAR motifs), and bHLH11dm-VP16 (VP16, an established activation domain) on bHLH105. Compared with the significant repression effect of bHLH11 on bHLH105, bHLH11dm had no significant effect, whereas bHLH11dm-VP16 enhanced the transactivation function of bHLH105 (Figure 6C). These data suggest that bHLH11 functions as a transcriptional repressor and that this function is dependent on its EAR motifs.

**Figure 6.**
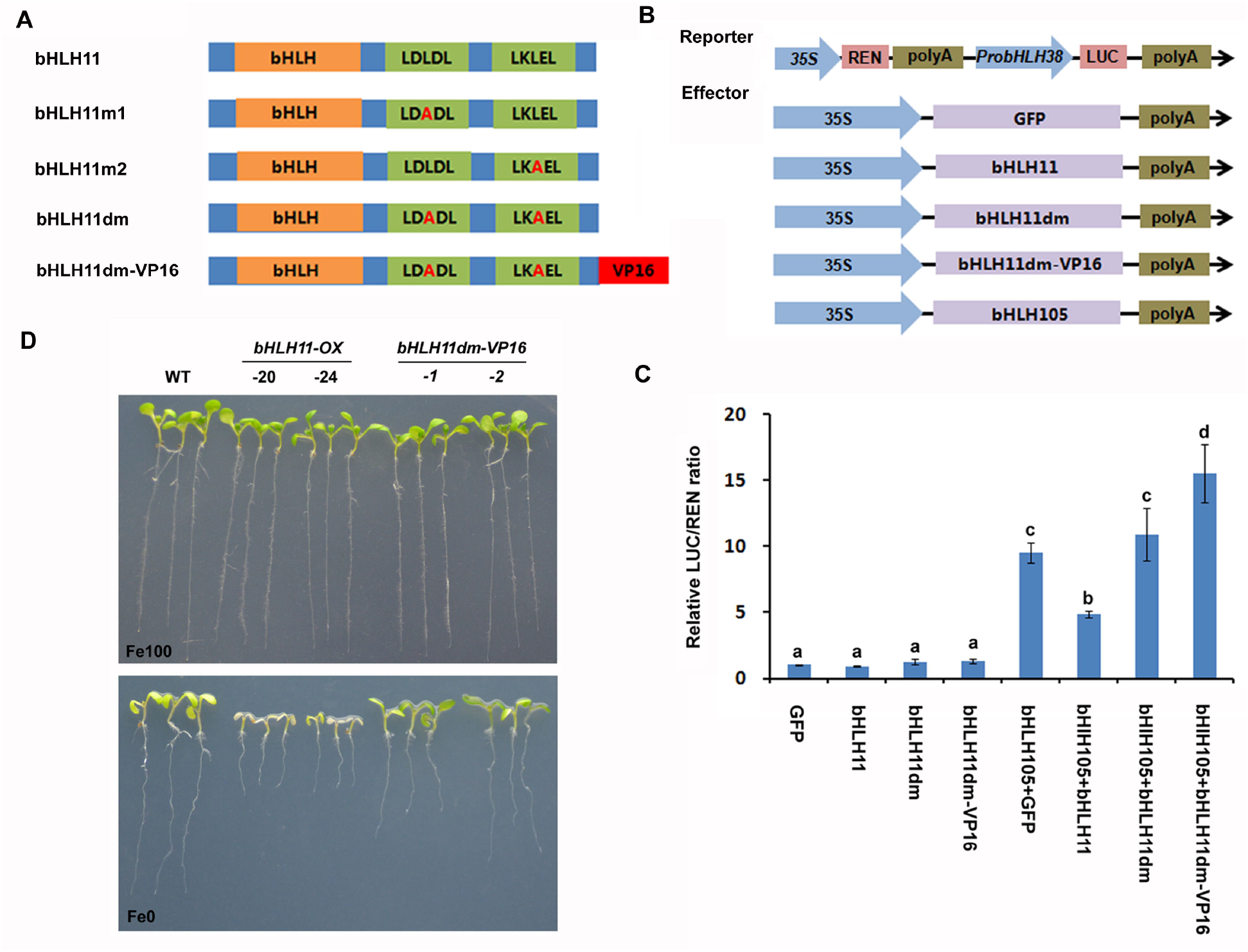
bHLH11 acts as a repressor dependently on its two EAR motifs. (A) Schematic diagram of the various mutated versions of bHLH11. The mutated amino acid is indicated in red. bHLH11m1, the first EAR mutated. bHLH11m2, the second EAR mutated. bHLH11dm, both double EARs mutated. bHLH11dm-VP16, bHLH11dm fused with the VP16 domain. (B) The schematic diagram shows the constructs used in the transient expression assays in (C). (C) The EAR motifs are required for the repression of bHLH11. Arabidopsis protoplasts were used for transient expression assays. The abundance of *GFP* was normalized to that of *NPTII.* The different letters above each bar indicate statistically significant differences as determined by one-way ANOVA followed by Tukey’s multiple comparison test (P < 0.05). (D) Phenotypes of *bHLH11dm-VP16* and *bHLH11-OX* plants. Seven-day-old seedlings grown on Fe0 or Fe100 medium are shown.

To assess the consequences of disrupting the repression functions of bHLH11 *in vivo*, we generated *bHLH11dm-VP16* transgenic plants. The *bHLH11dm-VP16* plants showed the enhanced tolerance to Fe deficiency compared to *bHLH11-OX* plants (Figure 6D). Correspondingly, the expression of *IRT1* and *FRO2* was activated in the *bHLH11dm-VP16* plants (Figure S6). Taken together, the EAR motifs are needed for the repression function of bHLH11.

### bHLH11 interacts with the transcription corepressors TPL/TPRs

The EAR motif is a characteristic of proteins interacting with the TPL/TPRs which function as transcription corepressors (Szemenyei et al., 2008; Pauwels et al., 2010; Causier et al., 2012). Thus, we determined whether bHLH11 interacts with TPL/TPRs. Yeast two-hybrid assays indicated that bHLH11 interacts with TPL/TPRs (Figure 7A). To further investigate whether the EAR motifs are required for the interaction, the various EAR-mutated versions, bHLH11m1, bHLH11m2, and bHLH11dm, were tested. The results indicated that the interaction between bHLH11 and TPL/TPRs was dependent on the EAR motifs, as the mutation of both EAR motifs abolished the interaction between bHLH11 and TPL/TPRs.

**Figure 7.**
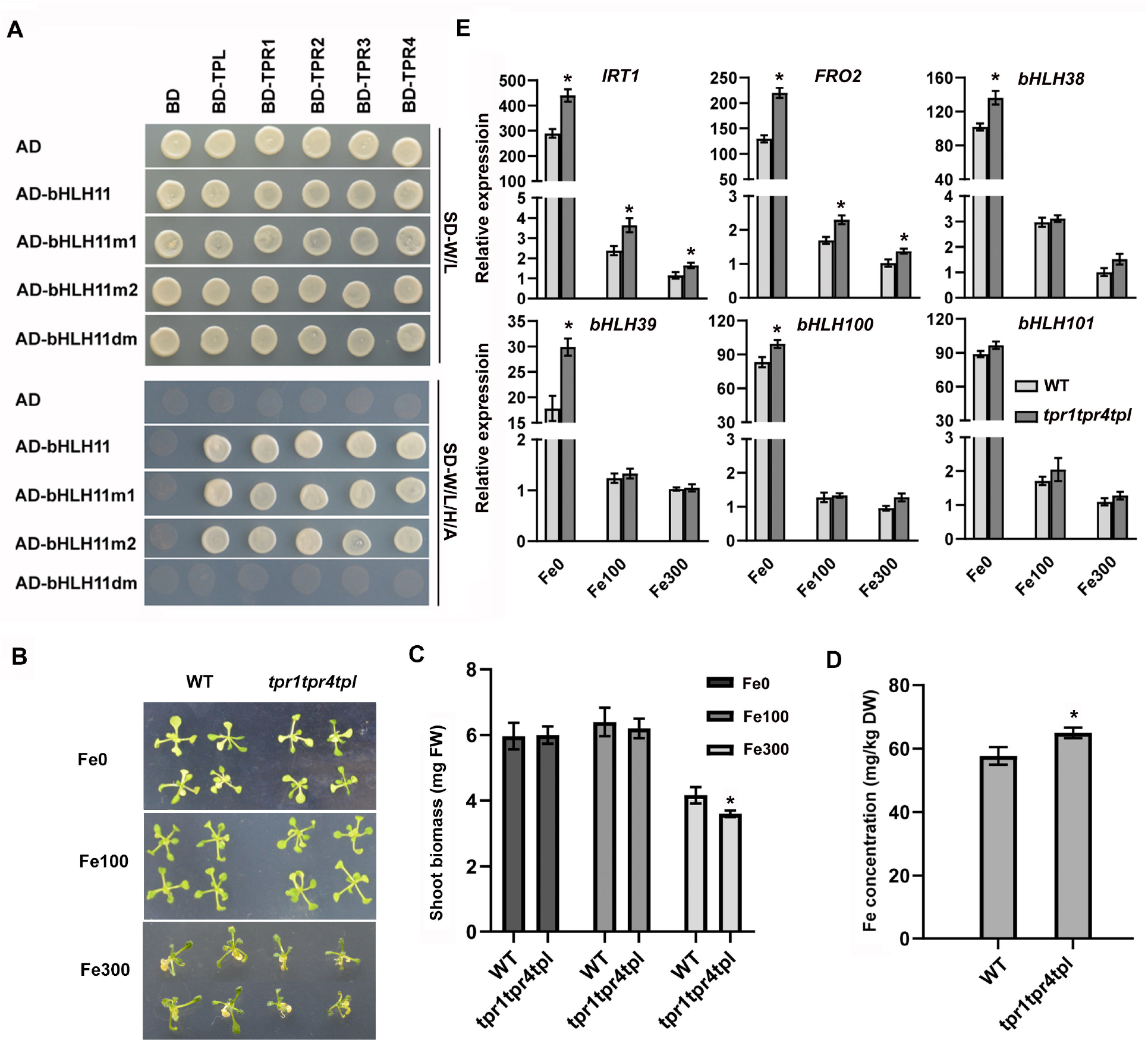
Enhanced Fe deficiency response in *tpr1 tpr4 tpl*. (A) The EAR motifs are required for the interaction between bHLH11 and TPL/TPRs. Yeast cotransformed with different BD and AD plasmid combinations was spotted on synthetic dropout medium lacking Leu/Trp (SD–T/L) or Trp/Leu/His/Ade (SD–T/L/H/A). (B) Phenotypes of *tpr1 tpr4 tpl*. Two-week-old seedlings grown on Fe0, Fe100 or Fe300 medium are shown. (C) Shoot biomass of *tpr1 tpr4 tpl*. Fresh weight of two-week-old shoots grown on Fe0, Fe100 or Fe300 medium. Three biological duplicates, each of which contains 15 plants, were analyzed. (D) Fe concentration of rosette leaves of 3-week-old wild type and *tpr1 tpr4 tpl* plants grown in soil. (E) Expression of *IRT1, FRO2* and bHLH Ib genes. Four-day-old plants grown on Fe100 medium were transferred to Fe0, Fe100 or Fe300 medium for three days, and root samples were harvested and used for RNA extraction and RT-qPCR. (C-D) The asterisks indicate that the values are significantly different from the corresponding wild type value by Student’s t Test (P < 0.05).

Next, we wanted to know whether TPL/TPRs participate in the regulation of Fe homeostasis. To this aim, the *tpr1 tpr4 tpl* triple mutant plants were used for phenotypic analysis. When grown on Fe0 or Fe100 medium, no visible difference was observed. In contrast, when grown on Fe300 medium, the shoot biomass of the *tpr1 tpr4 tpl* triple mutant plants was higher than that of wild type (Figure 7B, C). The measurement of Fe concentration showed that the *tpr1 tpr4 tpl* triple mutant plants had higher Fe concentration than wild type (Figure 7D). We also examined the expression of Fe deficiency responsive genes, finding that the expression of *IRT1* and *FRO2* was higher in the *tpr1 tpr4 tpl* than that in the wild type (Figure 7E). These data suggest that TPL/TPRs negatively regulate the expression of Fe uptake genes.

### Genetic relationship between *bHLH11* and *FIT*

Tanabe et al. (2019) reported that bHLH11 negatively regulates Fe uptake by repressing *FIT* transcription. To explore whether the function of bHLH11 depends on FIT, we conducted genetic analysis. The *bHLH11-OX-20* line was crossed into the *fit-2* mutant, and homozygous *bHLH11-OX-20/fit-2* plants were identified. When grown on Fe100 or Fe300 medium, no visible difference was observed among the wild type, *fit-2, bHLH20-OX-20* and *bHLH11-OX-20/fit-2* plants (Figure 8A; Figure 7A). However, when grown on Fe0 medium, the *bHLH11-OX-20/fit-2* plants were more sensitive to Fe deficiency than *fit-2* and *bHLH11-OX-20* plants, as shown by the shorter roots and bleached leaves (Figure 8A; Figure S7B). These data suggest that bHLH11 has roles independent of FIT.

**Figure 8.**
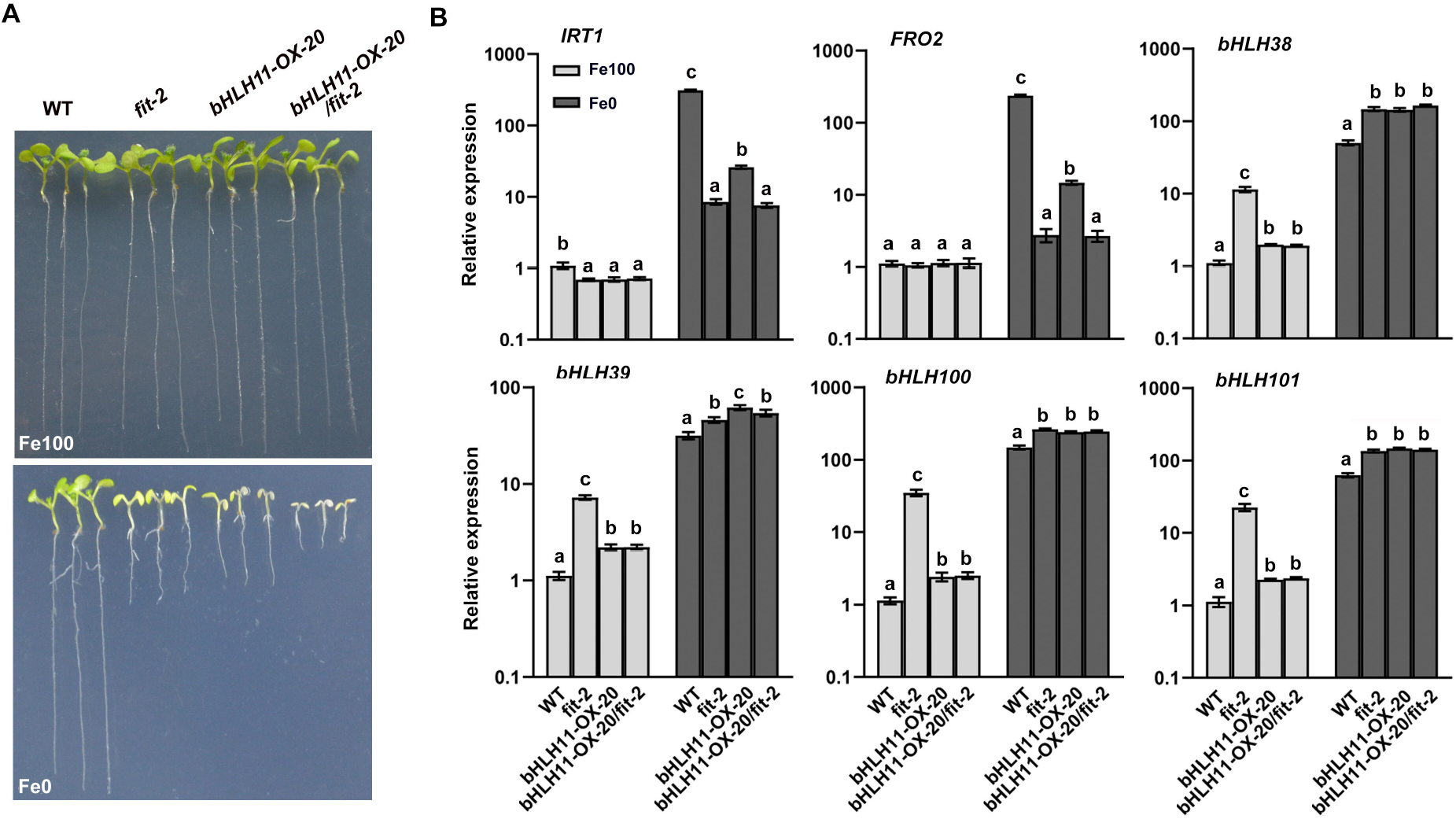
Genetic interaction between *bHLH11* and *FIT*. (A) Phenotypes of *bHLH11-OX-20/fit-2* plants. Seven-day-old seedlings grown on Fe0 or Fe100 medium are shown. (B) Expression of *IRT1, FRO2* and bHLH Ib genes. Four-day-old plants grown on Fe100 medium were transferred to Fe0 or Fe100 medium for three days, and root samples were harvested and used for RT-qPCR. The different letters above each bar indicate statistically significant differences as determined by one-way ANOVA followed by Tukey’s multiple comparison test (P < 0.05).

Subsequently, we examined the expression of *IRT1, FRO2* and bHLH Ib (Figure 8B). Under Fe deficient conditions, the expression of *IRT1* and *FRO2* was lower in the *fit-2* than that in the *bHLH11-OX-20*, but as low as that in the *bHLH11-OX-20/fit-2*. In contrast, the expression of bHLH Ib genes was similar among *fit-2, bHLH20-OX-20* and *bHLH11-OX-20/fit-2* under Fe deficient conditions. Under Fe sufficient conditions, although the expression of bHLH Ib genes was higher in the *fit-2*, the introduction of *bHLH11-OX-20* significantly repressed their expression. These data suggest that bHLH11 represses the expression of bHLH Ib genes independently of FIT under Fe sufficient conditions.

## Discussion

Plants sense Fe-deficient environments and activate a signal transduction cascade that ultimately results in the transcriptional regulation of downstream effector genes of the Fe uptake system. The expression of Fe homeostasis-associated genes is tightly regulated by Fe availability, including environmental Fe availability and local Fe availability in developing tissues and organs. However, this mechanism is not an on-off process but rather a fine-tuned one, with multiple layers of transcription regulations. Considerable progress has been made in deciphering the signal transduction pathways that maintain Fe homeostasis, leading to the identification of many signaling components. Here, we show that bHLH11 acts an active repressor by recruiting TPL/TPRs. bHLH11 contributes to Fe homeostasis by repressing bHLH IVc TFs.

The antagonistic regulation between positive and negative TFs is prevalent in plants. For example, the positive TFs MYC2/MYC3/MYC4 and the negative TFs bHLH3/bHLH13/bHLH14/bHLH17 antagonistically modulate jasmonic acid signaling (Fernandez-Calvo et al., 2011; Song et al., 2013). In Fe homeostasis signaling, the bHLH IVa TFs (bHLH18, bHLH19, bHLH20, and bHLH25) antagonize the bHLH Ib TFs to regulate FIT protein stability under Fe deficiency (Cui et al., 2018). We show the antagonistic function of bHLH11 to bHLH IVc TFs, which may explain why *bHLH11-OX* plants display the severe Fe deficiency phenotypes similar to those of bHLH IVc mutants (Liang et al., 2017). In addition to transcriptional regulation, the protein degradation is another type of regulation in Fe homeostasis signaling. As reported previously, bHLH105 and bHLH115 are degraded by BTS (Selote et al., 2015), and FIT by BTSL1 and BTSL2 (Rodríguez-Celma et al., 2019). We found that bHLH11 protein decreased under Fe deficient conditions (Figure 2B), which may benefit plants by alleviating the repression of bHLH11 to Fe uptake associated genes. Further investigation is required to understand the post-transcription regulation of *bHLH11* in response to Fe deficiency. These coordinated regulations of transcription and post-transcription may help plants adapt to their various Fe-nutrition habitats.

Although bHLH Ib genes were upregulated in both the *bHLH11-OX* and *bhlh11* mutant plants, we provide evidence supporting that bHLH11 negatively regulates bHLH Ib genes: (1) bHLH11 represses the promoter of *bHLH38* in the transient expression assays (Figure 5A); (2) the transient induction of *bHLH11* reduced the abundance of bHLH Ib genes in the *pER8-bHLH11* plants (Figure 5C); (3) the overexpression of *bHLH11* reduced the levels of bHLH Ib genes in the *fit-2* under Fe sufficient conditions (Figure 8B). We proposed that the upregulation of bHLH Ib genes may result from a feedback regulation that the severe Fe deficient status of *bHLH11-OX* plants activates bHLH Ib genes. In fact, feedback regulations are universal in Fe homeostasis. For example, bHLH Ib genes are also activated in the *irt1, frd3, opt3* and *fit* which are defective in Fe uptake or transportation (Wang et al., 2007). Therefore, our data suggest that the abundance of bHLH Ib genes are balanced by bHLH IVc TFs and bHLH11. We demonstrated that bHLH11 negatively regulates the expression of *IRT1* and *FRO2*, which is in consistence with the conclusion that bHLH11 represses the FIT-dependent Fe uptake (Tanabe et al., 2019). Meanwhile, bHLH11 also plays roles independent of FIT since its overexpression made *fit-2* more sensitive to Fe deficiency (Figure 8A), which is reasonable because bHLH11 can inhibit bHLH IVc TFs.

bHLH11 has no canonical NLS sequence. bHLH11 protein exists in the nucleus and cytoplasm, and it accumulates in the nucleus with the assistance of bHLH IVc TFs (Figure 4A). This interaction dependent nuclear localization of bHLH11 might contribute to the maintenance of Fe homeostasis. bHLH11 inhibits the activation ability of bHLH IVc TFs and restricts the expression of Fe uptake-associated genes. The repression function of bHLH11 may help plants avoid Fe toxicity and adapt to environments with excessive Fe by reducing the rate of Fe uptake. This putative protein shuttling strategy enables plants to respond quickly to Fe fluctuation.

Two types of transcriptional repressors exist: active and passive (Krogan and Long, 2009). Generally, active repressors repress transcription by recruiting transcriptional repression components, whereas passive repressors indirectly influence transcription by competitively interfering with activators. TPL/TPRs are a class of corepressors and usually recruited by the EAR motif containing transcriptional repressors (Kagale et al., 2010; Causier et al., 2012). bHLH11 functions negatively and has two EAR motifs, both of which can interact with TPL/TPRs, suggesting that bHLH11 is an active repressor. The observation that the EAR motif is conserved in bHLH11 homologs across different plant species, such as *Zea mays, Oryza sativa*, and *Brassica rapa* (Figure S8), implies that different plant species employ a conserved repression strategy to fine-tune Fe homeostasis. In Fe homeostasis signaling pathway, PYE is a negative regulator of Fe homeostasis associated genes *ZIF1, FRO3* and *NAS4* (Long et al., 2010), and an EAR motif was found in its C-terminal region (Kagale et al., 2010). ZAT12 (ZINC FINGER OF ARABIDOPSIS THALIANA 12) functions as a negative regulator of Fe uptake, and contains an EAR motif responsible for the interaction with FIT (Le et al., 2016). By contrast, EIN3 (ETHYLENE-INSENSITIVE 3), also containing an EAR motif (Kagale et al., 2010), is a positive regulator of Fe homeostasis by interacting with FIT (Lingam et al., 2011). It is likely that these TFs also recruit TPL/TPRs. In support of this hypothesis, the *tpr1 tpr4 tpl* plants are also sensitive to Fe excess to a lesser extent compared with the *bhlh11* plants, as shown by the shoot biomass (Figure 1B; Figure 7C).

This study expands our knowledge of the Fe homeostasis transcription network mediated by bHLH Ib and IVc proteins. Based on our findings, we propose a putative working model for bHLH11 (Figure 9). When Fe is sufficient, bHLH11 mRNAs increase and its proteins are stable. To limit Fe uptake, bHLH IVc proteins facilitate the accumulation of bHLH11 in the nucleus, where bHLH11 recruits TPL/TPR corepressors to repress the activation of bHLH IVc proteins to bHLH Ib genes, and then the reduction of bHLH Ib genes results in the down-regulation of Fe uptake genes *IRT1* and *FRO2.* When Fe is limited, bHLH11 proteins decrease rapidly and few bHLH11 proteins enter the nucleus to inhibit bHLH IVc. Finally, the bHLH Ib proteins accumulate and promote the expression of *IRT1* and *FRO2*. This enables plants to control Fe uptake and maintain Fe homeostasis. Our study provides experimental support for the existence of an elaborate system that allows plants to respond dynamically to Fe status. This mechanism is based on an equilibrium between the activation of Fe uptake-associated genes by bHLH IVc and their repression by bHLH11.

**Figure 9.**
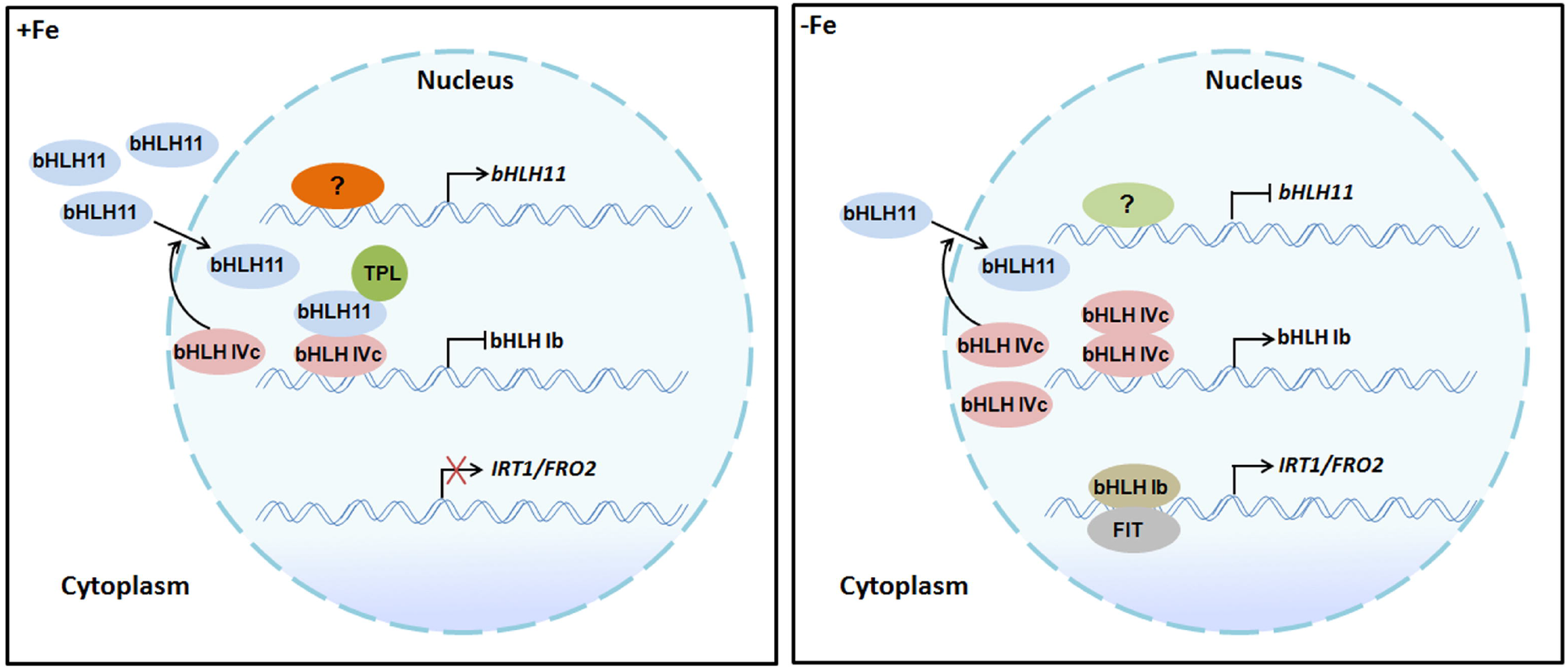
A working model of bHLH11 in Fe homeostasis. bHLH11 functions as an active repressor by recruiting TPL/TPR corepressors, and bHLH IVc TFs promote the nuclear accumulation of bHLH11. Under Fe-sufficient conditions, *bHLH11* message is activated and its protein accumulates. bHLH11 inhibits the transactivity of bHLH IVc TFs to bHLH Ib genes. The repression function of bHLH11 allows plants to avoid Fe toxicity. Under Fe-deficient conditions, unknown proteins repress the transcription of *bHLH11* and its protein decreases, which alleviates the bHLH11-mediated repression to bHLH IVc TFs. bHLH IVc TFs promote the transcription of bHLH Ib genes, and then bHLH Ib proteins interact with FIT to activate the expression of the Fe uptake genes *IRT1* and *FRO2*. The question signs indicate unknown proteins.

## Materials and methods

### Plant materials and growth conditions

*Arabidopsis thaliana* ecotype Col-0 was used as the wild type in this study. *fit-2* was described previously (Lei et al., 2020). *tpr1 tpr4 tpl* (N72353) was obtained from NASC. Plants were grown in long photoperiods (16-hour light/8-hour dark) or short photoperiods (10-hour light/14-hour dark) at 22°C. Surface sterilized seeds were stratified at 4°C for 2 d before being planted on medium. Half Murashige and Skoog (MS) medium with 1% sucrose, 0.8% agar A and the indicated Fe^2+^EDTA concentration were used. Fe0 (0 μM Fe^2+^), Fe50 (50 μM Fe^2+^), Fe100 (100 μM Fe^2+^) and Fe300 (300 μM Fe^2+^).

### Generation of CRISPR/Cas9-edited *bHLH11*

For CRISPR/Cas9-mediated editing of *bHLH11*, two target sites were designed by CRISPR-GE (Xie et al., 2017) to target the third and fourth exon of bHLH11, which were driven by the AtU3b promoter and respectively cloned into the pMH-SA binary vector carrying the Cas9 (Liang et al., 2016). The wild type plants were transformed and positive transgenic plants were selected on half-strength MS medium containing 20 μg/mL hygromycin. The positive transformants were sequenced and the homozygous mutants without the Cas9 were selected for further analysis.

### Generation of transgenic plants

HA-tag or VP16 domain were fused in frame with the full-length coding sequence of *bHLH11* to generate *35S:HA-bHLH11* and *35S:bHLH11dm-VP16* in the pOCA30 binary vector. *HA-bHLH11* was cloned into pER8 vector (Zuo et al., 2001). These constructs were introduced into *Agrobacterium tumefaciens* (EHA105) respectivley and then used for transformation in the wild type Arabidopsis. For complementation of *bhlh11-1,* the 3kb DNA fragment upstream of *bHLH11* translation start site was used to drive *HA-bHLH11-GFP* and then introduced into the *bhlh11-1* by *A. tumefaciens* mediated transformation.

### Yeast-two-hybrid assays

Full-length bHLH11 was cloned into pGBKT7 as a bait. The full-length of bHLH IVc in the pGADT7 was described previously (Li et al., 2016). Growth was determined as described in the Yeast Two-Hybrid System User Manual (Clontech).

### Subcellular localization

For the construction of *35S:bHLH11-mCherry*, mCherry-tag was fused with bHLH11. *35S:bHLH34-GFP, 35S:bHLH104-GFP, 35S:bHLH105-GFP, 35S:bHLH115-GFP*, and *35S:GFP* were described previously (Lei et al., 2020). *35S:bHLH11-mCherry* was co-expressed with various GFP-containing vectors in tobacco cells. Epidermal cells were recorded on an OLYMPUS confocal microscope. Excitation laser wave lengths of 488 nm and 563 nm were used for imaging GFP and mCherry signals, respectively.

### Fluorescence complementation assays

The tripartite split-GFP fluorescence complementation assay was described as previously (Lei et al., 2020). The C-end of bHLH11 was fused with the GFP11 tag. The N-end of bHLH IVc and bHLH121 was fused with the GFP10 tag. All vectors were introduced into *A. tumefaciens* (strain EHA105) and the various combinations of Agrobacterial cells were infiltrated into leaves of *Nicotiana benthamiana* by an infiltration buffer (0.2 mM acetosyringone, 10 mM MgCl_2_, and 10 mM MES, PH 5.6). Gene expression was induced 1 day after agroinfiltration by injecting 20 μM *β*-estradiol in the abaxial side of the leaves. Fluorescence of epidermal cells was recorded on a Carl Zeiss Microscopy.

### Co-immunoprecipitation assay

HA-bHLH11 and MYC-bHLH IVc or MYC-GFP were transiently expressed in the *N. benthamiana* leaves and the leaves were infiltrated with MG132 12 hours before harvesting. 2 g leaf samples were used for protein extraction in 2 ml IP buffer (50 m M Tris-HCl, pH 7.4, 150 mM NaCl, 1 mM MgCl_2_, 20% glycerol, 0.2% NP-40, 1 X protease inhibitor cocktail and 1 X phosphatase inhibitor cocktail from Roche). Lysates were clarified by centrifugation at 20, 000 g for 15 min at 4 °C and were immunoprecipitated using MYC antibody. IP proteins were analyzed by immunoblot using anti-HA and anti-MYC antibody respectively (Affinity Biosciences).

### Gene expression analysis

Total root RNA was extracted by the use of the Trizol reagent (Invitrogen). For the reverse transcription reaction, 1 μg total RNA was used for cDNA synthesis by oligo(dT)18 primer according to the manufacturer’s protocol (Takara). The resulting cDNA was subjected to relative quantitative PCR using the ChamQ™ SYBR qPCR Master Mix (Vazyme Biotech Co.,Ltd) on a Roche LightCycler 480 real-time PCR machine, according to the manufacturer’s instructions. For gene expression analysis in Arabidopsis plants, the relative level of genes was normalized to *ACT2* and *TUB2*. For the quantification of each gene, three biological replicates were used. The primers used for quantitative reverse transcription-PCR are listed in Table S1.

### Fe concentration measurement

To determine Fe concentration, rosette leaves from three-week-old seedlings grown in soil were harvested and dried at 65 °C for 3 days. About 100 mg dry weight was wet-ashed with 5 ml of 11 M HNO_3_ and 1 ml of 12 M HClO_4_ for 20 min at 220°C. Each sample was diluted to 16 ml with 18 MQ water and Fe concentration was analyzed on a Thermo SCIENTIFIC ICP-MS(iCAP6300).

### Transient expression assays in Arabidopsis protoplasts

Arabidopsis mesophyll protoplasts preparation and subsequent transfection were performed as described previously (Wu et al., 2009). The promoter sequence of *bHLH38* was amplified from genomic DNA and cloned into pGreenII 0800-LUC vector which contains a renillia luciferase encoding gene *REN* driven by the 35S promoter. The coding sequences of GFP and various kinds of bHLHs (bHLH34, bHLH104, bHLH105, bHLH115, bHLH11, bHLH11dm and bHLH11dm-VP16) were respectively cloned into the pGreenII 62-SK vector under control of 35S promoter. For the reporter and effectors, 10 μg plasmid for each construct was used. After protoplast preparation and subsequent transfection, firefly luciferase (LUC) and REN activities were measured using the Dual-Luciferase Reporter Assay System (Promega) following the manufacturer’s instructions. Relative (LUC) activity was calculated by normalizing against the REN activity.

### Transient expression assays in tobacco

*Agrobacterium tumefaciens* strains EHA105 was used in the transient expression experiments in tobacco. *pGAL4* promoter and BD domain were described previously (Li et al., 2016). *pGAL4* promoter was fused with NLS-GFP and cloned into the pOCA28 binary vector. 35S:BD, 35S:BD-bHLH104, 35S:BD-bHLH105 and 35S:HA-bHLH11 were constructed in the pOCA30 binary vector. For co-infiltration, different constructs were mixed prior to infiltration. Leaf infiltration was conducted in 3-week-old *N. benthamiana. NPTII* gene in the pOCA28 vector was used as the internal control. *GFP* transcript abundance was normalized to that of *NPTII*.

### Immunoblotting

For total protein extraction, roots were ground to a fine powder in liquid nitrogen and then resuspended and extracted in RIPA buffer (50 mM Tris, 150 mM NaCl, 1% NP-40, 0.5% sodium deoxycholate, 0.1% SDS, 1 mM PMSF, 1 x protease inhibitor cocktail [pH 8.0]). Isolation of cytoplasmic and nuclear proteins was performed as described previously (Li et al., 2018). Sample was loaded onto 12% SDS-PAGE gels and transferred to nitrocellulose membranes. The membrane was blocked with TBST (10 mM Tris-Cl, 150 mM NaCl, and 0.05% Tween 20, pH8.0) containing 5% nonfat milk (TBSTM) at room temperature for 60 min and incubated with primary antibody in TBSTM (overnight at 4°C). Membranes were washed with TBST (three times for 5 min each) and then incubated with the appropriate horseradish peroxidase-conjugated secondary antibodies in TBSTM at room temperature for 1.5 h. After washing three times, bound antibodies were visualized with ECL substrate.

## Supplemental data

**Supplemental Figure 1.** Identification of *bhlh11* mutants.

**Supplemental Figure 2.** Identification of *bHLH11* overexpression plants.

**Supplemental Figure 3.** Negative controls of tripartite split-sfGFP complementation assays.

**Supplemental Figure 4.** Antagonism between bHLH104 and bHLH11.

**Supplemental Figure 5.** Phenotypes of *pER8-bHLH11* transgenic plants.

**Supplemental Figure 6.** Expression of *IRT1* and *FRO2* in the *bHLH11dm-VP16* plants.

**Supplemental Figure 7.** Analysis of *bHLH11-OX-20/fit-2* plants.

**Supplemental Figure 8.** Conserved EAR motifs in the bHLH11 homologs from various plants species.

**Supplemental Table 1**. Primers used in this paper.

## Acknowledgements

We thank the Biogeochemical Laboratory and Central Laboratory (Xishuangbanna Tropical Botanical Garden) for assistance in the determination of metal contents. We also thank Germplasm Bank of Wild Species in Southwest China for confocal laser scanning microscopy.

## Finding

This work was supported by the National Natural Science Foundation of China (31770270) and the Applied Basic Research Project of Yunnan Province (2019FB028 and 202001AT070131).

## Author contributions

G.L. conceived the project. Y.L., R.L., M.P., C.L., Z. L., and G.L. constructed plasmids, M.P. and Y.C generated transgenic plants, and Y.L. and R.L. characterized plants, determined gene and protein expression, and conducted cellular assays. Y.L. and G.L. wrote the manuscript and all authors discussed and approved the manuscript.

## Conflict of interest statement

None declared

## Notes

### Competing Interest Statement

The authors have declared no competing interest.

